# Monkeypox Knowledge Graph: A comprehensive representation embedding chemical entities and associated biology of Monkeypox

**DOI:** 10.1101/2022.08.02.502453

**Authors:** Reagon Karki, Andrea Zaliani, Yojana Gadiya, Philip Gribbon

## Abstract

**Summary:** The outbreak of Monkeypox virus (MPXV) infection in May 2022 is declared a global health emergency by WHO. As of 28^th^ July, 21067 cases have been confirmed and the numbers are on the rise. Unfortunately, MPXV pathophysiology and its underlying mechanisms are not yet understood. Likewise, the knowledge of biochemicals and drugs used against MPXV and their downstream effects is sparse. In this work, using Knowledge Graph (KG) representations we have depicted chemical and biological aspects of MPXV. To achieve this, we have collected and rationally assembled several biological study results, assays, drug candidates, and preclinical evidence to form a dynamic and comprehensive network. The KG is compliant with FAIR annotations allowing seamless transformation and integration to/with other formats and infrastructures.

**Availability and implementation:** The Monkeypox knowledge graph is publicly available at https://github.com/Fraunhofer-ITMP/mpox-kg

**Supplementary information:** Supplementary data are available at Bioinformatics online.

## 1. Introduction

The recent COVID-19 pandemic has drastically changed the way research and scientific studies operate in areas of infectious and epidemic diseases. Although new discoveries and uncovering pathophysiology are the ultimate expectations, a new aspect that has been crucial to these is the response time (Khanna et al., 2020). Despite the expertise and technologies of the highest levels in hospitals, pharmaceutical companies, and research institutes, the response was not always timely. This clearly was the impact of lack of preparedness and since then we are determined to avoid experiencing the same chaos in future epidemics (Villa et al., 2020). One of the setbacks in such a situation was the unavailability of enough research data with metadata compliant with Findable, Accessible, Interoperable, and Reproducible (FAIR) data principles, consequently leading to mapping gaps between different domains of scientific studies. A number of efforts have emerged since then to harmonize sparse data and better understand the etiology of the disease (Harrison et al., 2021; Schmidt et al., 2021).

The ongoing multi-country outbreak of Monkeypox virus (MPXV) which started in May 2022 (https://www.ecdc.europa.eu/en/monkeypox-outbreak) has been declared a global health emergency and stands as another potential threat of pandemic. Unfortunately, the etiology of MPXV is not known and therefore, there is an urgent need to decipher it. This involves identifying the involvement of viral and host proteins in the infection, their interactions, virus replication biology, and potential drug candidates to perturb viral processes and mechanisms within the host. Additionally, from drug discovery and therapeutic perspective, it is important to know active molecules and their pharmacology either as a direct effect on viral-host interactions or as cellular toxicology effects. Understanding all the above-mentioned aspects will help accelerate drug repurposing and drug discovery processes. In this work, we have created a comprehensive Monkeypox Knowledge Graph (KG) that represents chemical-specific information such as chemicals and drugs active against MPXV along with their side effects, biological-specific information such as proteins, and their associated biological processes. The KG is represented with standard ontologies aligning it with the FAIR data principles. Furthermore, the KG is available in various graph formats to facilitate data handling and processing as required by the scientific community.

## 2. Material and methods

The overall methodology (Figure 1) used for the creation of the Monkeypox KG is divided into three main steps: i) biological/chemical resources identification ii) data harmonization and standardization iii) KG generation.

**Figure 1:**
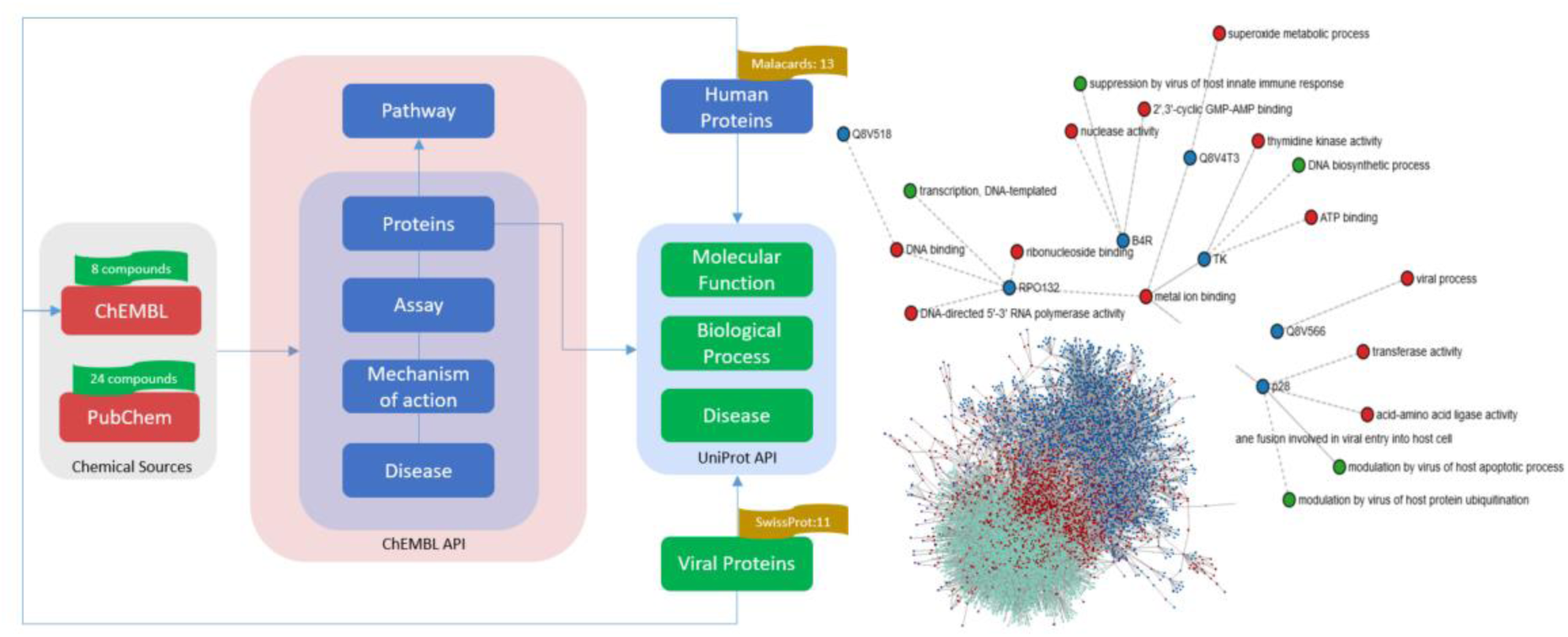
A schematic representation of the KG workflow (left) and visualization of the KG (right)

### 2.1 Resource identification

The chemical compounds used or tested against MPXV were retrieved from public chemical data resources, i.e. PubChem (Kim et al., 2021), ChEMBL (Davies et al., 2015) databases (last accessed: July 24, 2022). We queried PubChem with NCBI Taxonomy Identifier (ID) of the virus (NCBITaxon: 10244) and the associated chemicals were listed under the table “Chemicals and Bioactivities” and sub-table “Tested Compounds”. Since ChEMBL has its independent ontology for taxonomy, the database was queried using ChEMBL ID for MPXV (i.e., CHEMBL613120). Afterwards, we selected chemicals with pChEMBL value > 6 from either binding or functional assays since this condition ensures bioactivity of a given chemical. In general, this value is half-maximal response concentration/potency/affinity on a negative logarithmic scale. Next, using the taxonomy ID of MPXV, we collected reviewed protein entries (Swiss-Prot) from UniProt (Apweiler et al., 2004). For human proteins, we queried DISEASES, a human disease database, with DOID:3292 (Grissa et al., 2022). Lastly, Open Targets Platform was used to fetch information about “druggability” of proteins reported from studies (Ochoa et al., 2021).

### 2.2 Programmatic methods for data fetching and harmonization

The programmatic scripts and methods written were written in python (version 3.10) and are available at https://github.com/Fraunhofer-ITMP/mpox-kg. Firstly, we converted PubChem IDs to ChEMBL IDs because the information about chemical associated proteins, assays, mechanism of actions, pathways and diseases can be fetched from the ChEMBL API using ChEMBL IDs. After combining the identified proteins with proteins from UniProt and DISEASES, we used the UniProt API to extend the information on molecular functions, biological processes, sequences, pathways and diseases. Furthermore, we identified additional ChEMBL compounds that target proteins collected in the workflow and repeated the aforementioned steps of using ChEMBL and Uniprot API. Lastly, we mapped and harmonized chemical and protein names as they were collected from different resources and had different identifiers. For example, PubChem entry with compound ID 16124688 is registered in ChEMBL as CHEMBL1257073. We have used ChEMBL IDs for standard and uniform representation of chemicals. Similarly, Uniprot IDs were converted to HUGO names as it helps researchers to readily identify a given protein.

### 2.3 Construction of Monkeypox KG

The Monkeypox KG (Figure 1) is represented in the form of semantic triples using Biological Expression Language (BEL) with metadata annotation on nodes and relations using the PyBEL framework (Hoyt et al., 2018). PyBEL is a software tool built to facilitate data parsing, semantics validation, and visualization of data generated in BEL format. The framework provides a library of functions for exploring, querying and analyzing the KG. Moreover, the KG can be exported to other formats such as json, csv, sql, graphml and Neo4j which enables systematic comparison or integration with other KGs.

## 3. Results

While the literature and public data resources contain sparse MPXV related information, our approach has built a bridge between chemical and biological worlds, thus yielding a comprehensive KG. Starting from chemicals associated with MPXV, we were able to identify corresponding assays with bioactivity, target proteins, and their biological processes. Moreover, with MPXV and human proteins, we not only summoned knowledge about aforementioned aspects but also identified chemicals targeting these proteins.

Our query from PubChem retrieved 24 chemicals, all of which were successfully mapped to corresponding ChEMBL IDs. Similarly, 8 chemicals were retrieved from ChEMBL, out of which 6 were identical with the PubChem chemicals. The search in UniProt fetched 11 MPXV proteins whereas DISEASES returned 19 human proteins. Using these results as the primer for the KG, we created a KG using ChEMBL and UniProt API. The KG comprised 7216 nodes and 32676 relationships where we have identified 565 putative drugs targeting human and viral proteins. The full summary of KG statistics is available in the Supplementary File. The proteins represented in the KG were further labelled with “druggability” information using Open Targets. Additionally, we performed a sequence similarity search for the MPXV proteins and identified human homologues with sequence identity >35%. The results are also provided in Supplementary File.

## 4. Discussion

The COVID-19 pandemic has alerted the scientific community at different levels such as identifying early predictors, understanding pathogen biology and consequent pathophysiology, selecting first line of treatments, identifying putative drugs, and enabling data availability which is crucial for all sorts of research activities (Bibi et al., 2021; Domingo-Fernández et al., 2021). The lesson learned from COVID-19 is to be prepared and act immediately to avoid the next unprepared “COVID-19-esque” situation. In this regard, aligning with FAIR data principles, we have created a Monkeypox KG which is a comprehensive representation of biological and chemical entities associated with MPXV. To our knowledge, ours is the first KG in MPXV research.

One significant strength of our KG is that it not only embeds, harmonizes, and visualizes entities but also serves as a primer for downstream analyses. For example, a chemoinformatician can readily run similarity search analyses using the chemicals represented in the KG. Similarly, a biologist working with a certain protein and chemical can quickly find out other chemicals targeting the same protein. Considering these, we aim to facilitate ongoing and upcoming MPXV studies by serving a useful resource to different research groups and therefore will continue to update the KG actively. One of our next updates will include annotation of proteins with MPXV specific omics data. Finally, we plan to reach out to other public resources for enriching the knowledge in the KG.

## Authors’ contribution (eventually)

RK ideated and implemented the KG. RK, YG, AZ and PG wrote the paper.

## References

Apweiler, R., Bairoch, A., Wu, C. H., Barker, W. C., Boeckmann, B., Ferro, S., Gasteiger, E., Huang, H., Lopez, R., Magrane, M., & others. (2004). UniProt: the universal protein knowledgebase. Nucleic Acids Research, 32(suppl_1), D115–D119.

Bibi, N., Farid, A., Gul, S., Ali, J., Amin, F., Kalathiya, U., & Hupp, T. (2021). Drug repositioning against COVID-19: A first line treatment. Journal of Biomolecular Structure and Dynamics, 1–15.

Davies, M., Nowotka, M., Papadatos, G., Dedman, N., Gaulton, A., Atkinson, F., Bellis, L., & Overington, J. P. (2015). ChEMBL web services: streamlining access to drug discovery data and utilities. Nucleic Acids Research, 43(W1), W612–W620.

Domingo-Fernández, D., Baksi, S., Schultz, B., Gadiya, Y., Karki, R., Raschka, T., Ebeling, C., Hofmann-Apitius, M., & Kodamullil, A. T. (2021). COVID-19 Knowledge Graph: a computable, multi-modal, cause-and-effect knowledge model of COVID-19 pathophysiology. Bioinformatics, 37(9), 1332–1334.

Grissa, D., Junge, A., Oprea, T. I., & Jensen, L. J. (2022). DISEASES 2.0: a weekly updated database of disease–gene associations from text mining and data integration. Database, 2022.

Harrison, P. W., Lopez, R., Rahman, N., Allen, S. G., Aslam, R., Buso, N., Cummins, C., Fathy, Y., Felix, E., Glont, M., & others. (2021). The COVID-19 Data Portal: accelerating SARS-CoV-2 and COVID-19 research through rapid open access data sharing. Nucleic Acids Research, 49(W1), W619–W623.

Hoyt, C. T., Konotopez, A., & Ebeling, C. (2018). PyBEL: a computational framework for Biological Expression Language. Bioinformatics, 34(4), 703–704.

Khanna, R. C., Cicinelli, M. V., Gilbert, S. S., Honavar, S. G., & Murthy, G. V. S. (2020). COVID-19 pandemic: Lessons learned and future directions. Indian Journal of Ophthalmology, 68(5), 703.

Kim, S., Chen, J., Cheng, T., Gindulyte, A., He, J., He, S., Li, Q., Shoemaker, B. A., Thiessen, P. A., Yu, B., & others. (2021). PubChem in 2021: new data content and improved web interfaces. Nucleic Acids Research, 49(D1), D1388–D1395.

Ochoa, D., Hercules, A., Carmona, M., Suveges, D., Gonzalez-Uriarte, A., Malangone, C., Miranda, A., Fumis, L., Carvalho-Silva, D., Spitzer, M., & others. (2021). Open Targets Platform: supporting systematic drug–target identification and prioritisation. Nucleic Acids Research, 49(D1), D1302–D1310.

Schmidt, C. O., Fluck, J., Golebiewski, M., Grabenhenrich, L., Hahn, H., Kirsten, T., Klammt, S., Löbe, M., Sax, U., Thun, S., & others. (2021). Making COVID-19 research data more accessible-building a nationwide information infrastructure. Bundesgesundheitsblatt, Gesundheitsforschung, Gesundheitsschutz, 64(9), 1084–1092.

Villa, S., Lombardi, A., Mangioni, D., Bozzi, G., Bandera, A., Gori, A., & Raviglione, M. C. (2020). The COVID-19 pandemic preparedness or lack thereof: from China to Italy. Global Health & Medicine.

