## Supplementary File for "Monkeypox Knowledge Graph: A comprehensive representation embedding chemical entities and associated biology of Monkeypox"

### Outline

1. Monkeypox Knowledge Graph in numbers
2. Proteins from ‘druggability’ family
3. Viral-host protein predictions

**Supplementary Text**

1. **Knowledge Graph in numbers**

The Monkeypox KG (**Supplementary Figure 1)** is composed of 7216 nodes out of which 4226, 2195, 1427 and 387 nodes account for pathology, biological process, abundance and protein respectively. The pathology is further sub-divided to namespaces SideEffect and Disease where the corresponding numbers are 3319 and 907. Likewise, biological process is sub-divided to namespaces Gene Ontology Biological Process (GOBP), Gene Ontology Molecular Function (GOMF), Reactome and Mechanism of Action (MOA) and the corresponding numbers are 774, 537, 751 and 133. The abundance consisted of ChEMBL assays and chemicals where the numbers are 862 and 565 respectively. Lastly, there are 10 viral proteins and the rest 377 are human proteins (**Supplementary Figure 2**).

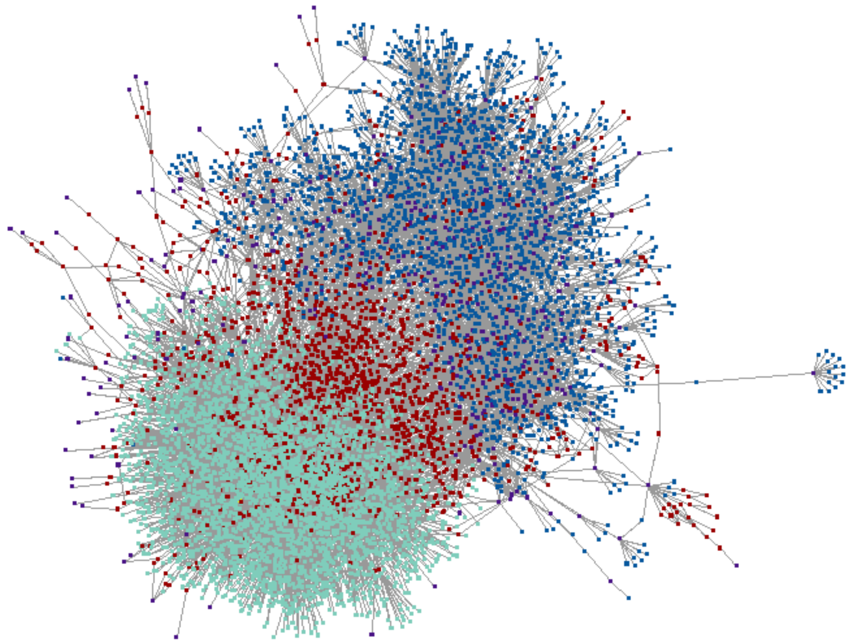

**Supplementary Figure 1**: A snapshot of the Monkeypox KG

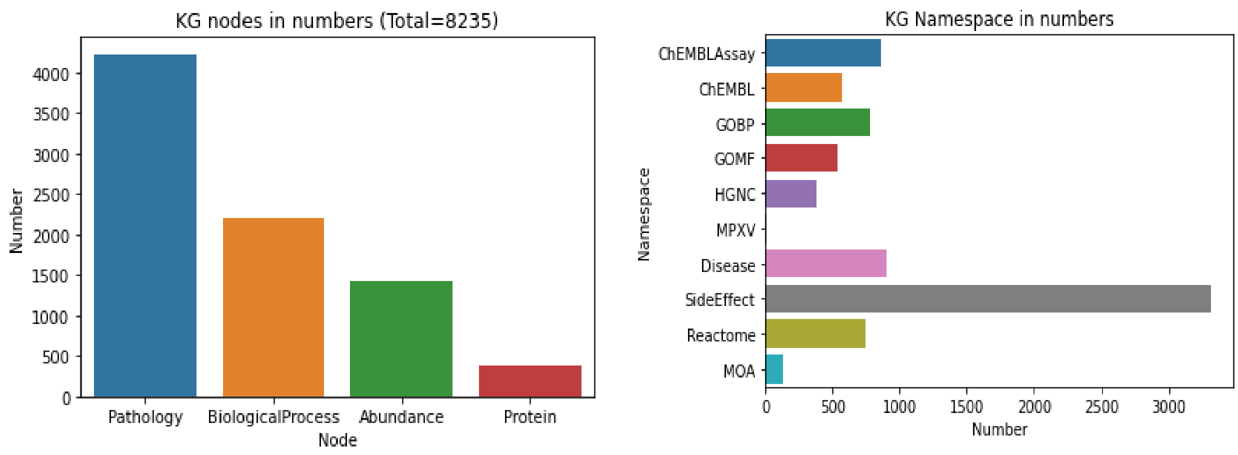


**Supplementary Figure 2**: Distribution of different nodes (left) and namespaces (right) in KG

#### 2. Proteins from "druggability" family

Using Open Targets, we have annotated the proteins in KG with the information whether they belong to the druggable family of proteins. We found out that 255 proteins out of 377 belonged to the druggable family of proteins (**Supplementary Table 1**).

| Protein | Druggable Family | Protein | Druggable Family | Protein | Druggable Family | Protein | Druggable Family |
|---|---|---|---|---|---|---|---|
| ABCB1 | Yes | CHRNB4 | Yes | HDAC2 | Yes | NR3C2 | Yes |
| ABCC8 | Yes | CNR1 | Yes | HDAC3 | Yes | OPRL1 | Yes |
| ACE | Yes | CNR2 | Yes | HDAC8 | Yes | OR10G6 | No |
| ACHE | Yes | COMT | Yes | HIF1A | Yes | OR51E2 | Yes |
| ADORA1 | Yes | CSNK2A1 | Yes | HPGD | Yes | P2RX1 | Yes |
| ADORA2B | Yes | CSNK2B | No | HRAS | Yes | P2RX3 | Yes |
| ADORA3 | Yes | CYB5R3 | No | HRH1 | Yes | P2RX4 | Yes |
| ADRA1A | Yes | CYP11B1 | Yes | HRH2 | Yes | P2RY2 | Yes |
| ADRA1B | Yes | CYP11B2 | Yes | HRH3 | Yes | PDE3A | Yes |
| ADRA1D | Yes | CYP17A1 | Yes | HRH4 | Yes | PDE4A | Yes |
| ADRA2A | Yes | CYP19A1 | Yes | HSD11B1 | Yes | PDE4B | Yes |
| ADRA2B | Yes | CYP1A1 | Yes | HSD11B2 | Yes | PDE5A | Yes |
| ADRA2C | Yes | CYP1A2 | Yes | HSD17B10 | Yes | PDE6H | Yes |
| ADRB1 | Yes | CYP1B1 | Yes | HSD17B3 | Yes | PGR | Yes |
| ADRB2 | Yes | CYP26A1 | Yes | HSP90AA1 | Yes | PKM | Yes |
| AGTR1 | Yes | CYP2B6 | Yes | HTR1A | Yes | PLG | Yes |

| | | | | | | | |
|---|---|---|---|---|---|---|---|
| AHCY | Yes | CYP2C19 | Yes | HTR1B | Yes | PMP22 | Yes |
| AHCYL1 | No | CYP2C9 | Yes | HTR1D | Yes | POLA1 | Yes |
| AKR1C1 | Yes | CYP2D6 | Yes | HTR2A | Yes | POLB | Yes |
| AKR1C2 | Yes | CYP3A4 | Yes | HTR2B | Yes | PPARG | Yes |
| AKR1C3 | Yes | CYP51A1 | Yes | HTR2C | Yes | PPAT | Yes |
| ALDH1A1 | Yes | DHFR | Yes | HTR3A | Yes | PPIA | Yes |
| ALOX15 | Yes | DRD1 | Yes | HTR4 | Yes | PPP3CA | Yes |
| ALOX5 | Yes | DRD2 | Yes | HTR5A | Yes | PRSS1 | Yes |
| ALOX5AP | Yes | DRD3 | Yes | HTR6 | Yes | PTGFR | Yes |
| AOC3 | Yes | DRD4 | Yes | HTR7 | Yes | PTGS1 | Yes |
| APEX1 | Yes | EBP | Yes | HTT | Yes | PTGS2 | Yes |
| AR | Yes | EGFR | Yes | IDO1 | Yes | RAB9A | Yes |
| ASAH1 | No | ERBB2 | Yes | IFNA1 | No | RECQL | Yes |
| ATP1A1 | Yes | ERBB4 | Yes | IMPDH1 | Yes | RGS17 | No |
| ATP4A | Yes | ESR1 | Yes | IMPDH2 | Yes | RIPK2 | Yes |
| BCHE | Yes | ESR2 | Yes | IRF3 | No | RXRA | Yes |
| BHMT | Yes | F12 | Yes | ITGAL | Yes | SCN1A | Yes |
| BLM | Yes | F2 | Yes | KCNH2 | Yes | SCNN1A | Yes |
| BMP2K | Yes | FABP3 | No | KCNJ11 | Yes | SERPINA6 | No |
| C1S | Yes | FABP4 | Yes | KCNJ8 | Yes | SHBG | Yes |
| C4A | No | FDPS | Yes | KCNN4 | Yes | SIGMAR1 | Yes |
| C4B | No | FFAR1 | Yes | KCNQ1 | Yes | SIRT1 | Yes |
| C5 | No | FGF1 | Yes | KDM4E | Yes | SLC12A1 | Yes |
| CA1 | Yes | FKBP1A | Yes | KDR | Yes | SLC12A3 | Yes |
| CA12 | Yes | FUT7 | No | KIT | Yes | SLC22A12 | Yes |
| CA14 | Yes | GAA | Yes | KLF5 | Yes | SLC22A6 | Yes |
| CA2 | Yes | GABBR1 | Yes | KLRC3 | No | SLC22A8 | Yes |
| CA3 | Yes | GABRA1 | Yes | KLRK1 | No | SLC29A1 | Yes |
| CA4 | Yes | GABRA2 | Yes | KMT2A | Yes | SLC6A2 | Yes |
| CA5B | Yes | GABRA3 | Yes | L3MBTL1 | Yes | SLC6A3 | Yes |
| CA6 | Yes | GABRA5 | Yes | LMNA | Yes | SLC6A4 | Yes |

| | | | | | | | |
|---|---|---|---|---|---|---|---|
| CA7 | Yes | GAK | Yes | MAOA | Yes | SLC7A11 | No |
| CA9 | Yes | GALE | Yes | MAOB | Yes | SLCO1B1 | Yes |
| CACNA1C | Yes | GFER | Yes | MAP2K1 | Yes | SMN1 | Yes |
| CACNA1G | Yes | GLO1 | Yes | MAPK1 | Yes | SQLE | Yes |
| CACNA1I | Yes | GMNN | Yes | MAPK10 | Yes | TACR1 | Yes |
| CBR1 | Yes | GPR35 | Yes | MAPK13 | Yes | TACR2 | Yes |
| CCL26 | No | GPR55 | Yes | MAPK14 | Yes | TACR3 | Yes |
| CCR4 | Yes | GRIA4 | Yes | MAPK9 | Yes | TDP1 | Yes |
| CD4 | No | GRIK1 | Yes | MAPT | Yes | THPO | Yes |
| CD46 | No | GRIK2 | Yes | MEN1 | No | THRA | Yes |
| CD55 | No | GRIK5 | Yes | METAP2 | Yes | THRB | Yes |
| CD8A | No | GRIN1 | Yes | MGLL | Yes | TMEM97 | No |
| CDK1 | Yes | GRIN2A | Yes | MKNK2 | Yes | TOP1 | Yes |
| CDK2 | Yes | GRK2 | Yes | MT-ND4 | Yes | TOP1MT | Yes |
| CDK4 | Yes | GRM1 | Yes | MTOR | Yes | TP53 | Yes |
| CDK5 | Yes | GRM4 | Yes | NAPRT | No | TPO | Yes |
| CES1 | Yes | GRM5 | Yes | NFKB1 | Yes | TRPA1 | Yes |
| CHRM1 | Yes | GSK3B | Yes | NISCH | Yes | TSHR | Yes |
| CHRM2 | Yes | GSTP1 | Yes | NPSR1 | Yes | TUBB4B | Yes |
| CHRM3 | Yes | HASPIN | Yes | NQO1 | Yes | TYMS | Yes |
| CHRM4 | Yes | HBB | Yes | NQO2 | Yes | UMPS | Yes |
| CHRNA1 | Yes | HDAC1 | Yes | NR1I2 | Yes | VKORC1 | Yes |
| CHRNB2 | Yes | HDAC11 | Yes | NR3C1 | Yes | WDR5 | Yes |

**Supplementary Table 1**: A table of proteins with corresponding druggability family information

#### 3. Viral-host protein predictions

*In silico* identification of possible viral-host protein interactions is a challenging task. In recent years, several attempts have been made to find structural similarity from sequence (Zhou et al., 2014). We have implemented the sequence-based approach taking 11 MPXV proteins and BLASTing them (Altschul et al., 1990) towards human sequence SWISSPROT collection (Bairoch & Apweiler, 2000). Only human homologues with sequence identity >35% and only four of MPXV proteins (i.e., Q8V571 (p28), P04363 (TK), Q8V4S4 (B4R), Q8V4Y0 (E8L)) have been found having a sequence similarity with such a minimal level. All four proteins, however, showed high similarity on relevant sequence lengths (in average ca. 200 residues). The list of the BLAST result is shown in **Supplementary Table 2**. The full BLAST result is available in https://github.com/Fraunhofer-ITMP/mpox-kg/tree/main/data/uniprot.

| Mpox Protein (Uniprot) | Mpox Protein Names | Human Protein (Uniprot) | Human Protein Names | Percent Identity |
|---|---|---|---|---|
| Q8V571.1 | p28 | Q13064.1 | MKRN3 | 47.143 |
| Q8V571.1 | p28 | Q13434.1 | MKRN4P | 48.333 |
| Q8V571.1 | p28 | Q9UHC7.3 | MKRN1 | 38.889 |
| Q8V571.1 | p28 | Q9H000.2 | MKRN2 | 38.095 |
| Q8V571.1 | p28 | O76064.1 | RNF8 | 41.304 |
| P04363.2 | TK | P04183.2 | TK1 | 67.836 |
| Q8V4S4.1 | B4R | Q8IYM2.2 | SLFN12 | 34.983 |
| Q8V4S4.1 | B4R | Q6IEE8.4 | SLFN12L | 35.644 |
| Q8V4Y0.1 | E8L | P07451.3 | CA3 | 37.281 |
| Q8V4Y0.1 | E8L | Q8N1Q1.1 | CA13 | 37.555 |
| Q8V4Y0.1 | E8L | P35218.1 | CA5A | 35.484 |
| Q8V4Y0.1 | E8L | P00915.2 | CA1 | 36.321 |
| Q8V4Y0.1 | E8L | P00918.2 | CA2 | 36.889 |

**Supplementary Table 2:** A table summarizing BLAST results for MXPV proteins

Using this list of viral and human proteins, we created a subgraph of the KG. We found out that processes ‘zinc ion binding’ and ‘carbonate dehydratase activity’ were shared between human proteins CA1, CA2, CA3 and viral protein Q8V4Y0. Additionally, we also identifed ‘metal ion binding’ as a shared process between viral proteins p28 and TK (Supplementary Figure 3).

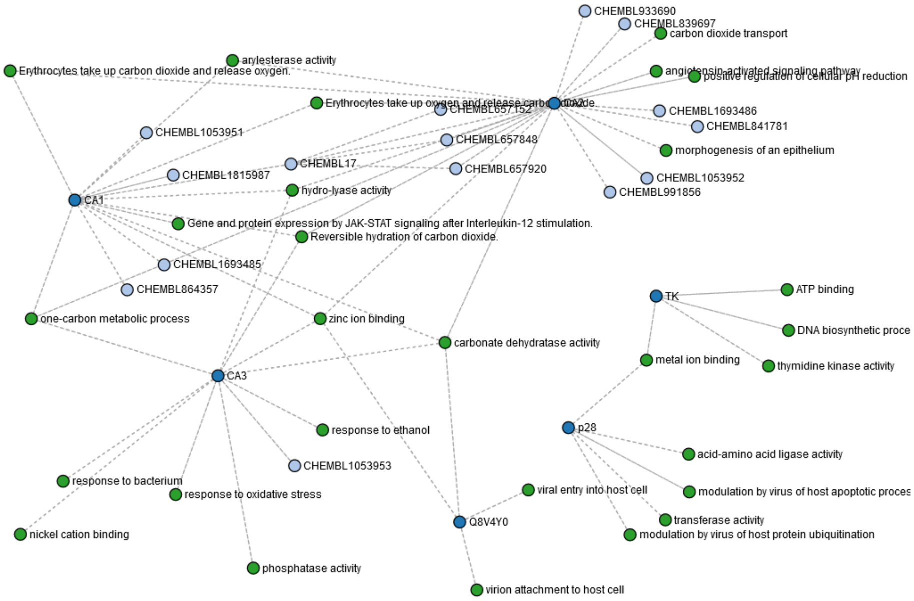


**Supplementary Figure 3**: A sub-graph depicting shared processes between human and viral proteins

#### References

Altschul, S. F., Gish, W., Miller, W., Myers, E. W., & Lipman, D. J. (1990). Basic local alignment search tool. *Journal of Molecular Biology*, *215*(3), 403–410.

Bairoch, A., & Apweiler, R. (2000). The SWISS-PROT protein sequence database and its supplement TrEMBL in 2000. *Nucleic Acids Research*, *28*(1), 45–48.

Zhou, H., Gao, S., Nguyen, N. N., Fan, M., Jin, J., Liu, B., Zhao, L., Xiong, G., Tan, M., Li, S., & others. (2014). Stringent homology-based prediction of H. sapiens-M. tuberculosis H37Rv protein-protein interactions. *Biology Direct*, *9*(1), 1–30.
